# Early-life adversity modulates growth trajectories and red blood cell mitochondrial metabolism in king penguin chicks

**DOI:** 10.1101/2025.08.22.671494

**Authors:** Nina Cossin-Sevrin, Katja Anttila, Mathilde Lejeune, Céline Bocquet, Maëlle Fusillier, Thomas Faulmann, Camille Lemonnier, Natacha Garcin, Anne Cillard, Pierre Bize, Jean-Patrice Robin, Suvi Ruuskanen, Vincent A Viblanc

**Author notes:** shared co-last authorship (Suvi Ruuskanen, Vincent A Viblanc). Corresponding author: Nina Cossin-Sevrin, Department of Biology, 20014 University of Turku, Finland. Institute of Biotechnology, HiLIFE, University of Helsinki, Finland.

## Abstract

In many avian species, variation in breeding phenology is known to affect reproductive success. In king penguins, the breeding cycle lasts more than a year, and the start of egg-laying extends over 3 months, with early-breeders laying eggs around December and late-breeders around February. Consequently, late-born chicks have less time to grow and build up their energy reserves before the Austral winter. It is however not known whether late-born chicks display alternative physiological strategies to catch up before winter. Variation in metabolic rate is one important pathway driving differences in growth patterns, as it is directly involved both in energy allocation processes and fitness. The conversion of resources into energy occurs in mitochondria, and the efficiency of this conversion is likely to play a fundamental role in explaining individual heterogeneity in growth and survival. The purposes of this study were to investigate the differences in red blood cell mitochondrial metabolism between early- and late-born king penguin chicks and to assess whether morphometric phenotypes could be explained by differences in mitochondrial metabolism. Late-chicks expressed higher mitochondrial metabolism compared to early-chicks at 100 days post-hatching, probably linked to the stress related to winter environmental conditions and a decrease in parental feeding rates. We did not find a clear association between chick mitochondrial metabolism and growth patterns, suggesting that mostly environmental conditions contributed in explaining different metabolic phenotypes. As king penguin populations in Crozet may face changing breeding conditions (changes in feeding area and foraging trips duration) in relation to global changes, studying the physiological traits and adaptations underlying the chick growth and survival may help understanding the response of king penguins facing challenging conditions.

## Introduction

Two important aspects influence the success of the breeding season: the timing of breeding (i.e. phenology) and the investment of individuals in breeding - both influencing reproductive performance and fitness (Both et al., 2006; Gilsenan et al., 2020; Shipley et al., 2020; Verhulst & Nilsson, 2008). On one hand, the timing of breeding depends on environmental cues, such as photoperiod and ambient temperature, both of which influence the peak availability of food resources in the environment (Bonamour et al., 2019; Helm et al., 2013; Marrot et al., 2018; Nilsson & Källander, 2006; Visser et al., 1998). On the other hand, the timing of breeding is also influenced by breeder experience (age) and body condition (e.g. physiological capacity for the female to produce eggs). In many species, experienced and older breeders tend to breed earlier in the season (e.g. blackbirds, kittiwakes) (Coulson & White, 1956; Perrins, 1970; Snow, 1958). Phenological variation during the breeding season has been extensively studied to understand the effect of such variation on reproductive success (review in Dunn, 2004). Yet, this remains a pressing issue, as several studies raised advancement in phenology (for both migration and reproduction) in response to global changes in many species (Both et al., 2006; Charmantier & Gienapp, 2014; Dunn, 2004; Dunn et al., 2010; Hällfors et al., 2020).

According to the match-mismatch hypothesis the timing of breeding and subsequent offspring growth should be synchronized with the peak of resource availability to maximize the breeding success (Both et al., 2006; Kharouba & Wolkovich, 2020; Ramírez et al., 2017; Stenseth et al., 2002). Breeding early in the season (but not too early) can bring several advantages, including more chances to find a good breeding habitat (e.g. nest, cavity, territory in a colony), to match the maximal food availability period, but also more chance to find a good partner (Arvidsson & Neergaard, 1991; Both et al., 2006; Coté, 2000; Currie et al., 2000; Sutton & Freeman, 2023). For some species, early-born offspring experience lower predation rate compared to late-born ones (Coté, 2000; Götmark, 2002; Sutton & Freeman, 2023). All these reasons contribute to explain why experienced breeders tend to breed early in the season in many species, and why selection pressures often favor early breeding (De Villemereuil et al., 2020; Marrot et al., 2018; Van Noordwijk et al., 1995; Visser et al., 1998).

Among avian species with a large variation in the timing of reproduction, king penguins (*Aptenodytes patagonicus*) have a laying onset distributed over several months during the austral summer (November to mid April): with early-breeders (egg hatching around January) and late-breeders (egg hatching around March). As the single chick reaches independence about 12 months after hatching, a successful breeding season (including pre-breeding molt) takes more than one year to complete (Brodin & Olsson, 1998; Stonehouse, 1956, 1960; Viblanc et al., 2014; Weimerskirch et al., 1992). During winter, king penguin chicks remain in the colony on land (gathered all together in crèches), while parental food provisioning decreases in relation to the changes in the antarctic polar front position (main feeding area during breeding) (Bost et al., 1997; Saraux et al., 2012; Scheffer et al., 2010). Therefore, chicks may experience prolonged fasting periods during winter, leading to significant body mass loss (usually *ca* 40%) and a high mortality, especially for late-born chicks (Bost et al., 2015; Fernandes, 2023; Stier et al., 2014; VanHeezik et al., 1993; Verrier, 2003).

Successful breeders attempting to breed the following season will be delayed and start breeding later in the austral summer. For the late-born chicks (hereinafter late-born *vs.* early-born chicks are referred to as late-chicks *vs*. early-chick), the growth period is drastically reduced, resulting in a limited amount of time to accumulate body lipids and protein content reserves before winter, and as consequence late-chicks rarely survive the winter (first winter survival probability: 0.64 *vs*. 0.29 for early- and late-chicks respectively, Fernandes et al., 2025; Fernandes, 2023; Stier et al., 2014; VanHeezik et al., 1993). Prior studies showed that late-chicks can have a higher body mass and faster growth rates than early-chicks at the beginning of the growth period (Stier et al., 2014; VanHeezik et al., 1993). However, this difference is not maintained over the entire growth period: late-chicks are smaller and lighter than early-chicks before winter, and remain smaller at fledging (i.e. departure at sea) (Fernandes, 2023; Stier et al., 2014; VanHeezik et al., 1993; Verrier, 2003). Beside different growth trajectories, early- and late-chicks also differ in their physiology: late-chicks have higher blood corticosterone levels, higher oxidative stress and DNA damage, but also have reduced blood cell telomeres length compared to early-chicks at 10 days post-hatching (Stier et al., 2014). Despite short-term benefits, accelerated growth rates most likely come at a cost for the king penguin chicks (Geiger et al., 2012). Overall, a late breeding attempt in the season has consequences not only on the offspring survival and breeding success, but also on the offspring phenotype.

On one hand, variation in growth trajectories and survival between early- and late-chicks could be linked to environmental factors, including forecast variability, access to nutritional resources (parental feeding rates) or predation pressure (with small chicks more likely predated) (Descamps et al., 2005; Eichhorn et al., 2011). One the other hand, variation in growth and survival could be linked to physiological determinants differing between the two groups of chicks. Among plastic phenotypic traits, variation in metabolic rate is one important pathway driving variation in growth patterns, as it is directly involved both in energy allocation processes, and in individual fitness (Brown et al., 2018; Burger et al., 2019, 2021; Norin & Metcalfe, 2019). The conversion of resources into energy occurs through oxidative phosphorylation in mitochondria, where metabolic fuels are transformed into energy (ATP synthesis). The efficiency of this conversion, as well as the rate of production of pro-ageing compounds is likely to play a fundamental role in explaining individual heterogeneity in growth and survival (Goodrich & Clark, 2023; Heine & Hood, 2020; Koch et al., 2021; Salin et al., 2019). Previous research work investigated the modulation of whole-organism metabolic rate, but also the modulation of skeletal muscle mitochondrial respiration in king penguin chicks in response to winter-acclimation and fasting (Bourguignon et al., 2017; Duchamp et al., 1989; Monternier et al., 2014; Teulier et al., 2013), though whether mitochondrial metabolism differs depending on the timing at which early- and late-chicks were born remains to be determined. Such variation may be expected since the food resources fueling mitochondrial function, and the regularity and amount of provisioning patterns, are known to differ (in mammals and humans: Kyriazis et al., 2022; in birds: Cooper-Mullin et al., 2021; in fish: Salin et al., 2016).

The purposes of this study were to investigate i) the differences in growth trajectories and ii) red blood cell (RBC) mitochondrial metabolism between early- and late-born king penguin chicks. As avian RBCs are nucleated, mitochondrial metabolism can be measured using blood sample and used as a proxy of respiration in other tissues, enabling less-invasive and longitudinal studies (Casagrande et al., 2023; Cossin-Sevrin, 2024; Koch et al., 2021; Stier et al., 2017; Thoral et al., 2024b). Although RBCs may not directly be involved in the chick growth (as other tissues and organs), their function of carrying oxygen to the whole-organism is expected to affect the individual growth. Moreover, prior studies demonstrated RBC mitochondrial metabolism to correlate with the whole-animal metabolism (Koch et al. 2021; Casagrande et al. 2023; Thoral et al. 2024a), and with other tissues (e.g. muscle in king penguins, Stier et al., 2017). We aimed to assess if potential differences in growth patterns between early-and late-chicks could be linked to a modulation of RBC mitochondrial metabolism and its efficiency during the whole growth period. King penguin chicks experience a rapid body mass increase during their first summer (i.e. hereafter refers to core growth period), and then (usually) a decrease in body mass during winter when parental feeding decreases (Fernandes, 2023; Stier et al., 2014; VanHeezik et al., 1993; Verrier, 2003). To monitor these different stages, we collected blood samples in early- and late-chicks at different time-points: during the core growth period (35 days post-hatching), at the end of the core growth period (100 days post-hatching), and during winter (150 days post-hatching). As prior literature showed that late-chicks express higher markers of oxidative stress (Stier et al, 2014), we expected late-chicks to express higher RBC mitochondrial metabolism (potentially leading to a higher production of oxidative stress markers). For example, higher oxidative stress in late-chicks could be linked to differences in mitochondrial proton leak, or in the usage of mitochondrial complex I, which is known to be a contributor to cellular production of reactive oxygen species (ROS) (Hirst, 2013; Wirth et al., 2016). Higher mitochondrial metabolism would coincide as well with the energy requirements linked to a faster growth rate for the late-chicks. Because a decrease in blood cell mitochondrial content with the age during the growth period has been reported in several species (e.g. great tits, japanese quails, Cossin-Sevrin et al., 2022, 2023; Stier et al., 2022) we expected RBC mitochondrial metabolism of king penguin chicks to decrease with the age of the chicks.

## Material and Methods

### Ecology of king penguins and study area

This study was conducted on the Possession Island in the Crozet Archipelago (46°25′S; 51°52′E) during the breeding seasons 2020-2021 and 2021-2022 in the king penguin colony “La Grande Manchotière, Baie du Marin”, hosting ca 20,000 breeding pairs (Barbraud et al., 2020). Following courtship and egg-laying, the single chick is fed by both parents until the beginning of winter. During winter, the chick remains on land in crèches in the colony and experiences long periods with a few feeding events, or no feeding at all (Cherel et al., 1987; Saraux et al., 2012; Weimerskirch et al., 1992). Parents start by regularly feeding the chick again during spring until the chick reaches independence a few months later (beginning of summer): when the chick completed its juvenile molt and is able to feed itself, i.e. more than *ca* 12 months after egg laying (Ancel et al., 2013; Stonehouse, 1960; Weimerskirch et al., 1992). As a result, king penguin breeders that had a successful breeding season cannot complete their breeding molt and start the following breeding cycle on time. The onset of egg-laying is distributed over several months, with early breeding attempts between November / December (hatching around January) and late breeding attempts between January / February (hatching around March) (Fig.1).

**Fig.1:**
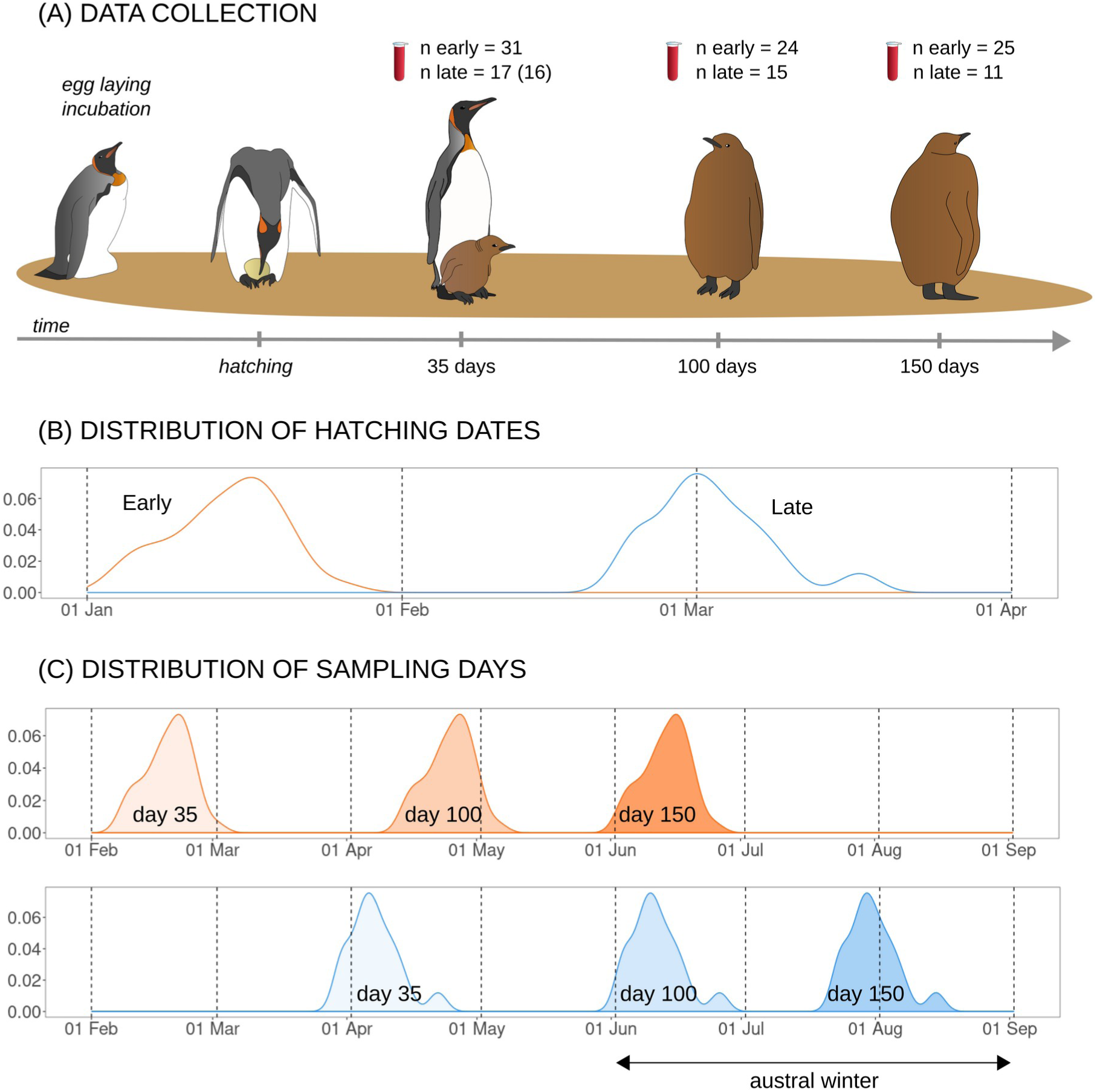
Schema of the study design. Data collection of RBC mitochondrial metabolic rates measurements (A), density of the distribution of hatching dates (B), density of the distribution of sampling days (C) across the breeding season for early-*vs.* late-chicks (C). Note that the scale differs for (B) and (C). Sample-sizes for RBC mitochondrial respiration measurements are indicated. A total of 55 individuals were included in this study: N_early_ = 31 and N_late_ = 23 individuals. For 35 days measurements, one value was missing for *ROUTINE* (sample size in brackets). Final sample-sizes were lower than expected due to missing data (e.g. logistical constraints such as unfavorable weather conditions and restricted access to the colony).

Data were collected on both early-born (N_2020_ = 16, N_2021_ = 16 individuals) and late-born king penguin chicks (N_2020_ = 15, N_2021_ = 8 individuals). Breeding pairs were randomly selected in the border of the colony (to minimize disturbance) during courtship and marked with a non-permanent animal dye (Porcimark, Kruuse, Langeskov, Denmark). Breeding pairs were monitored daily to record egg-laying and hatching dates (± 24h). The day where the chick was found completely outside of the egg shell was counted as “day 1”. As the parents alternate between foraging trips at sea and fasting periods on land in the colony to complete incubation and brooding tasks, we closely monitored the identity of the parent (female or male) present on land with the offspring. Females and males were visually sexed according to the order of parental alternation in the colony (males are the first to incubate), vocalization and morphometrics (Kriesell et al., 2018; Weimerskirch et al., 1992). Chicks included in this study were marked with a fishtag (placed at 20 days post-hatching), and a Darvic plastic flipper-band from 100 days post-hatching. Darvic flipper-bands were removed before the completion of the juvenile molt and the first departure at sea (*ca* 325 days post-hatching).

To investigate differences in growth trajectories, metabolism and survival between early- and late-chicks, king penguin chicks were captured in the colony for morphometric measurements and blood sampling (see below) at 35 days (mean ± SD = 35.2 ± 0.5 days), 100 days (99.9 ± 0.6 days) and 150 days (149.5 ± 1.2 days) post-hatching (see Fig.1). At 35-days post-hatching, the chick is experiencing a period of rapid body mass gain, ending towards 100 days post-hatching. In the beginning of winter (respectively *ca* 150 post-hatching days for early-chicks and 100 days for the late-chicks in this study), the chick is experiencing a reduced number of feeding events by the parents (Geiger et al., 2012; Saraux et al., 2012; Stier et al., 2014). According to the situation, the capture procedure was as follows. (i) If the chick was incubated by the parent during the procedure, we covered the adult’s eyes with a hood to reduce handling-stress and provided a dummy egg to the adult while delicately removing the chick. The chick was carefully placed in a warm bag using a hot water bottle until the end of the procedure and then replaced under the parental brood patch. (ii) If the chick was not incubated by the parent, the parent (if present) usually moved away over a few meters upon experimenter approach, leaving the chick behind. The chick was then captured and returned to the colony in the same area after the procedure. The return of the adult was then monitored from a distance, using binoculars, until the chick was reunited with its parent. Chick body mass was obtained using a spring scale (± 5g) for 35 day-old chicks and using an electronic scale (± 2g) for 100 and 150 days-old chicks. Flipper length (an index of structural size) was measured using a solid metal ruler (± 1mm).

### RBC mitochondrial aerobic metabolism

To measure RBC mitochondrial aerobic metabolism, blood samples were collected from the marginal flipper vein using G25 needle fitted to a 1mL heparinized syringe for the 35 days-old chicks (maximum 0.9mL) and with a G23 needle fitted to a 2.5mL heparinized syringe for the 100 and 150 days-old chicks (maximum 1.5mL). Blood samples were immediately transferred (within *ca* 5-10 min) to a laboratory facility located in the vicinity of the king penguin colony for processing. After refrigerated centrifugation (+4°C, 5min, 800g), RBCs were resuspended in PBS and kept at +4°C until high-resolution respirometry measurements could be performed at the research station located a few kilometers from the colony, a few hours later (2 Oroboros Instruments, Innsbruck, Austria). RBCs were then centrifuged a second time to remove PBS (+4°C, 5min, 800g) and resuspended in MIR05 respiration buffer to measure mitochondrial metabolism on permeabilized RBC at 38°C using a similar protocol as described in Stier et al. (2019) and Cossin-Sevrin et al. (2023, 2025): digitonin (20µg/mL), pyruvate (5mM), malate (2mM), ADP (1.25mM), succinate (10mM), oligomycin (2.5µM), antimycin A (2.5 µM).

Five distinct respiration rates were calculated and analyzed: 1) the endogenous cellular respiration rate before permeabilization (*ROUTINE*), 2) the maximum respiration rate fueled with exogenous substrates of complex I (pyruvate/malate), as well as ADP (*CI*), 3) the maximum respiration rate fueled with exogenous substrates of complexes I and II (succinate), as well as ADP (*CI+II*), 4) the respiration rate contributing to the proton leak after adding oligomycin in the presence of exogenous substrates of complexes I and II (*LEAK*), 5) the respiration rate supporting ATP synthesis through oxidative phosphorylation (*OXPHOS_CI+II,_* calculated as *CI+II - LEAK*). Non-mitochondrial respiration was measured after the addition of antimycin A (2.5µM) (raw data average ± SD: 0.90 ± 0.30 pmol.s^-1^) and was subtracted from all respiration rates. We also calculated 2 mitochondrial flux ratios (FCR): 1) *OXPHOS* coupling efficiency (*OxCE,* calculated as *OXPHOS_CI+II_ / (OXPHOS_CI+II_ + LEAK)*, and 2) the proportion of maximal respiration capacity being used under endogenous cellular condition (*FCR_R/OL_)* calculated as *ROUTINE / (OXPHOS_CI+II_ + LEAK)*. Respiration rates were standardized by the quantification of total protein measured in each sample, using Pierce™ BCA Protein Assay Kit. Repeatability of the quantification of total protein (based on sample-duplicate) was: R [CI 95%] = 0.980 [0.976, 0.984]. Technical repeatabilities of mitochondrial metabolic rates were: *ROUTINE:* R [CI 95%] = 0.712 [0.451, 0.905]; CI: R = 0.937 [0.867, 0.98]; *CI+II*: R = 0.878 [0.733, 0.963]; *LEAK*: R = 0.841 [0.674, 0.949]; *OXPHOS_CI+II_*: R = 0.915 [0.816, 0.975] based on 9 duplicates.

### Statistical analysis

Statistical analyses were conducted using R v.4.0.2. software (https://www.r-project.org/). For morphometrics, our final sample-size included 137 measurements from 32 early-chicks and 23 late-chicks: n_early_ = 32, 27, 28 *vs.* n_late_ = 22, 16, 12 at day 35, 100 and 150 respectively. For RBC mitochondrial metabolism measurements, the final sample-size encompassed 123 measurements from 31 early-chicks and 17 late-chicks (see Figure 1 for additional information on sample sizes). Our final sample-size was lower than expected as a result of missing data (because of logistical constraints, such as bad weather and limited access to the colony).

To investigate the differences in growth trajectories between early- and late-chicks, we used linear mixed models (LMMs, *lme4* package in R) with the body mass, body size (flipper length) and body condition (scale mass index as calculated in Peig & Green, 2009) as response variables and included as fixed factors the age (3 levels: 35, 100 and 150 days post-hatching) in interaction with the hatching group (2 levels: early *vs.* late), and the year of sampling (2 levels: 2020 *vs.* 2021). Bird ID was included as random effect to take into account the non-independence of measures collected on the same chick.

To test if RBC mitochondrial metabolic rates were different between early- and late-chicks, we performed similar LMMs as described above (except for *LEAK*), using mitochondrial metabolic rates as response variables. Because of convergence issues, *LEAK* was analyzed using a generalized linear mixed model (GLMM, *lme4* package in R) with a gamma error distribution. We preliminary tested the interaction between the age and the group in the models, but removed it if non-significant in order to properly interpret the main effects without scaling variables. To complete LMMs, we also conducted linear discriminant analysis (LDA) using the *MASS* package in R (Venables & Ripley, 2002), where the hatching group of chicks (early *vs.* late) was discriminated according to RBC mitochondrial metabolic rates. With this complementary analysis, we aimed to test if we could separate early- and late-chicks according to their RBC mitochondrial metabolic profile, i.e. including *ROUTINE*, *CI, CI+II* and *LEAK. OXPHOS_CI+II_* and FCR(s) could not be included in the LDA because of their non-independence with other metabolic rates (e.g. *OXPHOS_CI+II_* calculated as *CI+II - LEAK*). To test if RBC mitochondrial metabolism was associated with the chick body mass, we performed separate LMMs using chick body mass as continous variable, and by including the age (3 levels: day 35, day 100, day 150), and year of sampling (2 levels: 2020 *vs.* 2021) as fixed effects. Bird ID was included as random effect. Following our findings (see results), we used complementary LMs to test if RBC mitochondrial metabolism measured at 100 days post-hatching were associated with the environmental context. To this aim, we used each RBC mitochondrial metabolic rates as a response variable and included the date of sampling (in number of weeks in the year) and the year of sampling (2 levels: 2020 *vs.* 2021) as fixed factors.

To assess the association between growth trajectories and RBC mitochondrial metabolism, we used LMs testing if body mass gain during the core growth period. To this aim, body mass gain (differences in body mass between 35 and 100 days post-hatching) was used as response variable. For each RBC mitochondrial metabolic rate, measurements collected at day 35 and day 100 were both included as continuous variables in the models, while the hatching group (2 levels: early *vs.* late) and the year of sampling (2 levels: 2020 *vs.* 2021) were included as fixed factors. Finally, to test the association between body mass change during winter and RBC mitochondrial metabolism, we performed similar analyses as described above, using the body mass changes between 100 and 150 days post-hatching as response variable.

Fledging success of the chicks was monitored (individuals fledgling around *ca* 325 days post-hatching), and was estimated using GLMs with logistic binary distribution of the response variable (survival: 0 = dead, 1 = alive), with the hatching group (2 levels: early *vs.* late) and year of sampling (2 levels: 2020 *vs.* 2021) included as fixed factors.

## Results

### Differences in growth trajectories and survival between early- and late-chicks

We found a significant interaction of the age and the hatching group for chick body mass (age*group: F_2.89.1_ = 61.68, P < 0.001), body size (age*group: F_2,78.8_ = 86.12, P < 0.001) and body condition (age*group: F_2,87.2_ = 19.8, P < 0.001). Body mass, size and condition did not differ between early- and late-chicks at day 35 (Tukey HSD *post hoc* comparisons: early *vs.* late, all P > 0.15). At 100 and 150 days post-hatching, however, late-chicks were lighter (Tukey HSD *post hoc* comparisons: early *vs.* late, day 100: -40.3% and day 150: -43.3% for late-chicks; all P < 0.001, see Table 1A, Fig.2A), smaller (Tukey HSD *post hoc* comparisons: early *vs.* late, day 100: -22.3% and day 150: -19.3% for late-chicks, all P < 0.001, see Table 1B, Fig.2B) and with lower body condition than early-chicks (Tukey HSD *post hoc* comparisons: early *vs.* late, day 100: -16.5% and day 150: -25.3% for late-chicks, all P ≤ 0.001; Table 1). Chick body condition was significantly lower (-10%) for the breeding season 2021-2022 (F_1.51.41_ = 11.49, P = 0.001). Fledging success (i.e. chick departure at sea) was significantly lower for late-chicks (early: 12 chicks fledged out of 32; late: 2 chicks fledged out of 23) (P = 0.01), but was not affected by the year of sampling (P = 0.26).

**Table 1:** Results of LMMs testing the effect of the interaction between the chick age (number of days post-hatching) and hatching group (early *vs*. late) on (A) chick body mass (g) and (B) chick body size (flipper length, mm). Observations day 35: n_early_ / n_late_ = 32/22; day 100: n_early_ / n_late_ = 27/16, day 150: n_early_ = 28/12. N=55 individuals in total. Final sample-sizes are lower than expected due to missing data (see Methods). Estimated marginal means are reported with their standard error. *Post hoc* comparisons results with Tukey HSD correction are presented for the early and late groups. Bird ID was included as random intercepts in models. σ2, within-group variance; τ00, between-group variance Sample size (n) along with marginal (fixed effects only) and conditional (fixed and random effects). Bold indicates significance (P<0.05).

| Predictors | A. Body mass (g) |  |  | B. Body size (flipper length, mm) |  |  |
| --- | --- | --- | --- | --- | --- | --- |
|  | Estimate | SE | P | Estimate | SE | P |
| (Intercept) | 3281.5 | 211.06 | **< 0.001** | 156.84 | 4.09 | **< 0.001** |
| age (day 100) | 4918.4 | 191.58 | **< 0.001** | 142.30 | 2.80 | **< 0.001** |
| age (day 150) | 4286.82 | 189.23 | **< 0.001** | 144.22 | 2.76 | **< 0.001** |
| hatching group (late) | -149.21 | 275.71 | 0.59 | -8.84 | 5.24 | 0.10 |
| year (2021) | -370.82 | 234.97 | 0.12 | 1.94 | 4.78 | 0.69 |
| age (day 100) x group (late) | -3080.96 | 314.33 | **< 0.001** | -57.99 | 4.70 | **< 0.001** |
| age (day 150) x group (late) | -3045.54 | 355.64 | **< 0.001** | -49.46 | 5.04 | **< 0.001** |
| *Post hoc* comparisons for: |  |  |  |  |  |  |
| day 35: early vs. late | 149 | 276 | 0.590 | 8.84 | 5.24 | 0.10 |
| day 100: early vs. late | 3230 | 311 | **< 0.001** | 66.82 | 5.66 | **< 0.001** |
| day 150: early vs. late | 3195 | 331 | **< 0.001** | 58.29 | 5.92 | **< 0.001** |
| Random effects: |  |  |  |  |  |  |
| $\sigma^2$ | 719.8 | | | 10.44 | | |
| $\tau_{00}$ bird | 682.4 | | | 15.59 | | |
| n observations | 137 |  |  | 136 |  |  |
| N individuals | 55 |  |  | 55 |  |  |
| Marginal $R^2$ /conditional $R^2$ | 0.91<br>/0.82 | | | 0.98<br>/0.93 | | |

**Fig.2:**
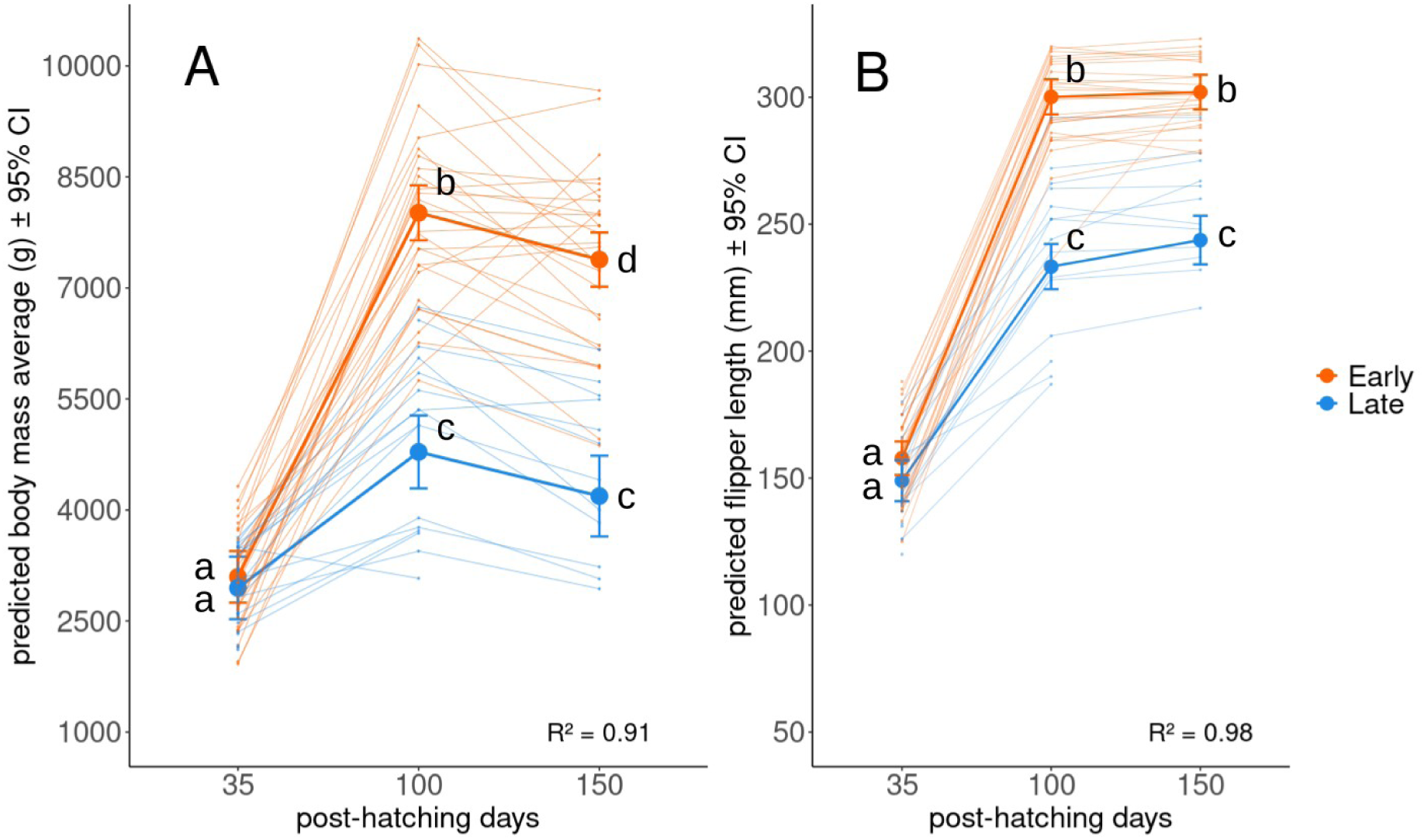
Predicted chick body mass (g) (A) and predicted chick body size (flipper length, mm) (B) from day 35 to day 150 post-hatching according to hatching group (early *vs.* late). Predicted averages with their 95% confidence interval (CI) and results from Tukey HSD post hoc tests are reported. The letters indicate significance of the *post-hoc* comparisons: if groups share a letter, they are not significantly different. R^2^ from LMMs are shown (see details and sample-sizes in Table 1). Raw data are plotted behind predicted average (lines). Orange color refers to the early-chicks, blue color refers to the late-chicks.

### RBC mitochondrial metabolism

The interaction between age and hatching group was statistically significant for *ROUTINE* (age*group: F_2,85.8_ = 6.96, P = 0.002) and *FCR_R/OL_* (age*group: F_2,84.7_ = 7.42, P = 0.001). For both *ROUTINE* and *FCR_R/OL_,* differences between early- and late-chicks were only significant at 100 days post-hatching, with higher RBC metabolic rates for late-chicks (Tukey HSD post-hoc comparisons early *vs.* late at day 100: all P < 0.001, Fig.3).

**Fig.3:**
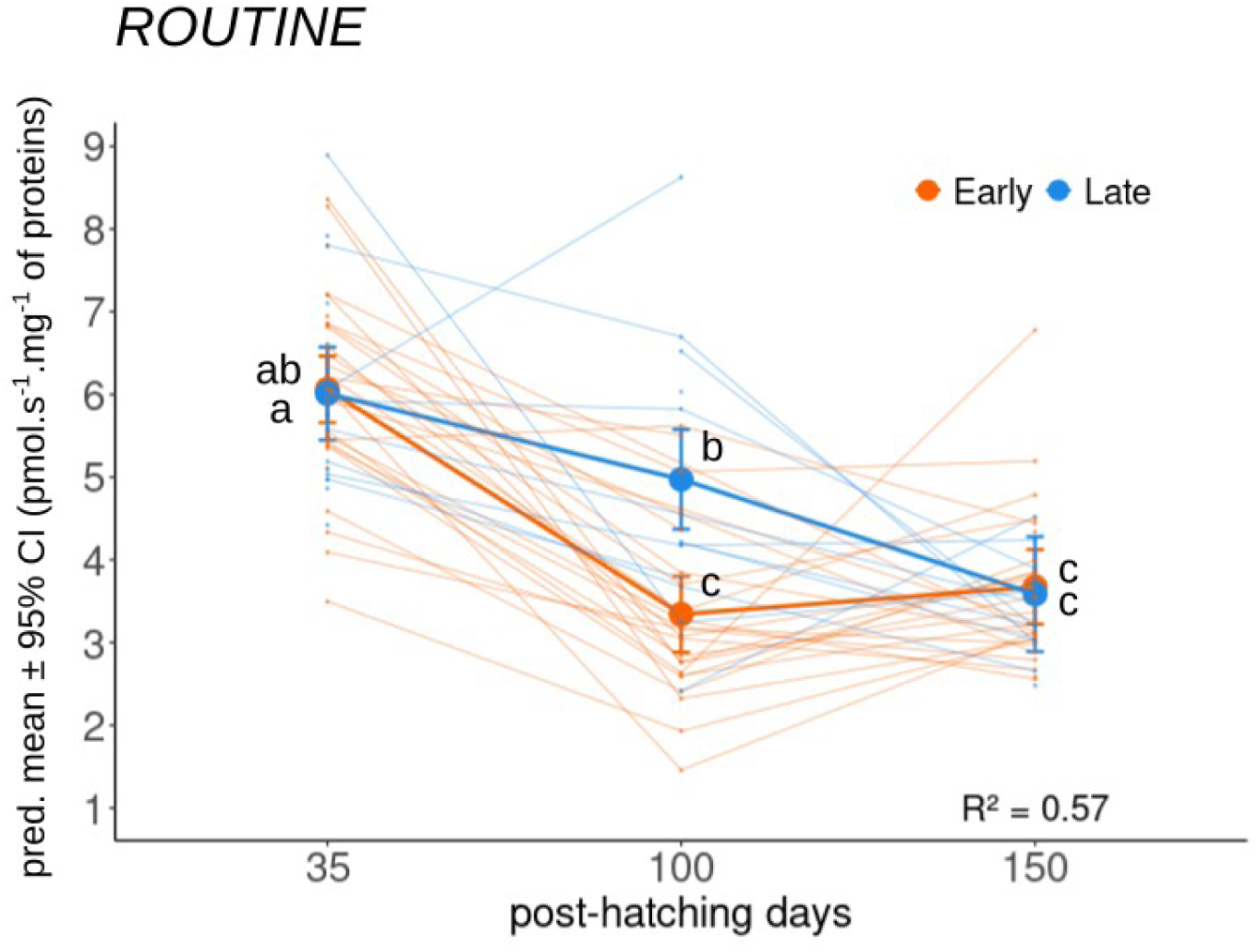
Predicted *ROUTINE* average (pmol.s^-1^.mg^-1^ of proteins) from day 35 to day 150 post-hatching according to hatching group (early *vs.* late). Predicted averages with their 95% confidence interval (CI) and results from Tukey HSD post hoc tests are shown. The letters indicate significance of the *post-hoc* comparisons: if groups share a letter, they are not significantly different. R^2^ from LMM is presented. Raw data are plotted behind predicted average (lines). Orange color refers to the early-chicks, blue color refers to the late-chicks.

C*I* was significantly higher in late-chicks than early-chicks (estimate ± SE = 0.63 ± 0.30, F_1,38.1_ = 4.32, P = 0.04, Table 2). In contrast, *CI+II, LEAK, OXPHOS_CI+II_* and *OXPHOS* coupling efficiency were not different between early- and late-chicks (Table 2).

**Table 2:** Results of Linear mixed models (and Generalized linear mixed model for *LEAK*) testing the effect of chick age (in number of days post-hatching), hatching group (early *vs.* late) and year (2020 *vs.* 2021) on chick RBC mitochondrial metabolic rates (pmol.s^-1^.mg^-1^ of proteins) and FCRs. Observations day 35: n_early_ / n_late_ = 31/17; day 100: n_early_ / n_late_ = 24/15, day 150: n_early_ = 25/11. N=55 individuals in total. *OxCE* refers to *OXPHOS* coupling efficiency. Estimates are reported with their standard error. σ2, within-group variance; τ00, between-group variance. Sample size (n) along with marginal (fixed effects only) and conditional (fixed and random effects). Bold indicates significance (P < 0.05).

| Predictors | *CI* |  | *CI+II* |  | *LEAK* |  | *OXPHOS<sub>CI+II</sub>* |  | *OxCE* |  |
| --- | --- | --- | --- | --- | --- | --- | --- | --- | --- | --- |
| | Estimate ± SE | P | Estimate ± SE | P | Estimate ± SE | P | Estimate ± SE | P | Estimate ± SE | P |
| (Intercept) | 5.96 ± 0.29 | **< 0.001** | 12.58 ± 0.40 | **< 0.001** | 0.59 ± 0.06 | **< 0.001** | 10.72 ± 0.37 | **< 0.001** | 201.66 ± 21.32 | **< 0.001** |
| age (day 100) | -1.84 ± 0.31 | **< 0.001** | -4.09 ± 0.43 | **< 0.001** | -0.32 ± 0.05 | **< 0.001** | -3.53 ± 0.39 | **< 0.001** | -0.02 ± 0.01 | **0.03** |
| age (day 150) | -2.23 ± 0.32 | **< 0.001** | -5.10 ± 0.45 | **< 0.001** | -0.41 ± 0.05 | **< 0.001** | -4.38 ± 0.40 | **< 0.001** | -0.03 ± 0.01 | **0.01** |
| hatching group (late) | 0.62 ± 0.30 | **0.04** | 0.54 ± 0.41 | 0.19 | 0.03 ± 0.06 | 0.58 | 0.43 ± 0.39 | 0.27 | -5$\times 10^{-4}$ ± 0.01 | 0.96 |
| year (2021) | -0.29 ± 0.29 | 0.33 | -0.16 ± 0.40 | 0.69 | 0.44 ± 0.06 | **< 0.001** | -0.99 ± 0.38 | **0.01** | -0.10 ± 0.01 | **< 0.001** |
| Random effects |  |  |  |  |  |  |  |  |  |  |
| $\sigma^2$ | 1.42 | | 1.99 | | 0.23 | | 1.77 | | 0.05 | |
| $\tau_{00}$ bird | 0.39 | | 0.41 | | 0.10 | | 0.61 | | 0.02 | |
| n observations | 123 |  | 123 |  | 123 |  | 123 |  | 123 |  |
| N individuals | 55 |  | 55 |  | 55 |  | 55 |  | 55 |  |
| Marginal R<sup>2</sup> /conditional R<sup>2</sup> | 0.39 /0.35 |  | 0.58 /0.57 |  | 0.51 /0.42 |  | 0.59 /0.54 |  | 0.56 /0.50 |  |

When conducting supplementary analyses to assess differences observed at day 100: *ROUTINE, CI* and *FCR_R/OL_* were not associated with chick body mass measured at day 100 days (all F < 0.88, all P > 0.35, Table S1, Fig.S1), but significantly increased with sampling date (number of weeks in the year) (all F > 5.12, all P < 0.03, Table S1, Fig.S1).

Except for *CI*, all RBC metabolic rates were significantly impacted by sampling year, with higher *ROUTINE*, *FCR_R/OL_* and *LEAK* (all F > 7.25, all P < 0.01), but lower *OXPHOS_CI+II_* and *OXPHOS* coupling efficiency in the breeding season 2021-2022 (year 2021) than 2020-2021 (year 2020) (all F > 6.77, all P < 0.01).

All RBC mitochondrial metabolic rates decreased with age (all F > 29.32, all P < 0.001, Table 2).

**Fig.4:**
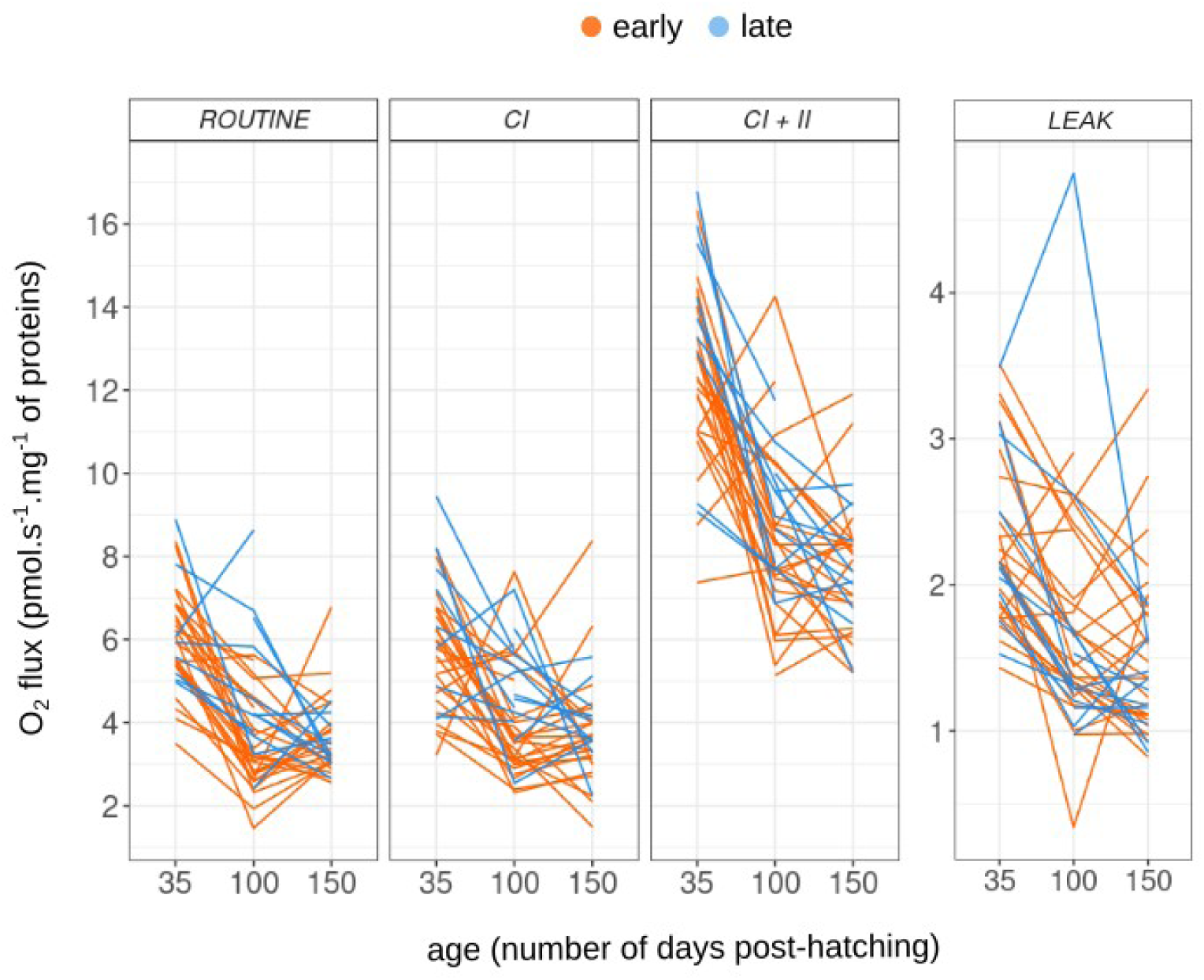
Changes in chick RBC mitochondrial metabolic rates (pmol.s^-1^.mg^-1^ of proteins) from day 35 to day 150 post-hatching according to hatching group (early *vs.* late). Raw data are presented. Orange color refers to the early-chicks, blue color refers to the late-chicks. Observations day 35: n_early_ / n_late_ = 31/17; day 100: n_early_ / n_late_ = 24/15, day 150: n_early_ = 25/11. N=55 individuals in total. See details of the models in Tables S1 and S2.

We used a linear discriminant analysis (LDA) to test the differences in RBC mitochondrial metabolic profile (i.e. *ROUTINE, CI, CI+II, LEAK*) between early- and late-chicks. Early and late-chicks were statistically different at day 100 (ANOVA: F_1,37_ = 15.49, P < 0.001, Fig.5), but not at 35 days or 150 days post-hatching (ANOVA: all F < 2.19, P > 0.15, Fig.5). When comparing RBC mitochondrial metabolic profile measured in early- and late-chicks during a similar period during the season (i.e. early-chicks at day 150 and late-chicks at day 100, see Fig.1), LDA analysis revealed a significant difference between groups (ANOVA: F_1,38_ = 12.32, P = 0.001, Fig.5).

**Fig.5:**
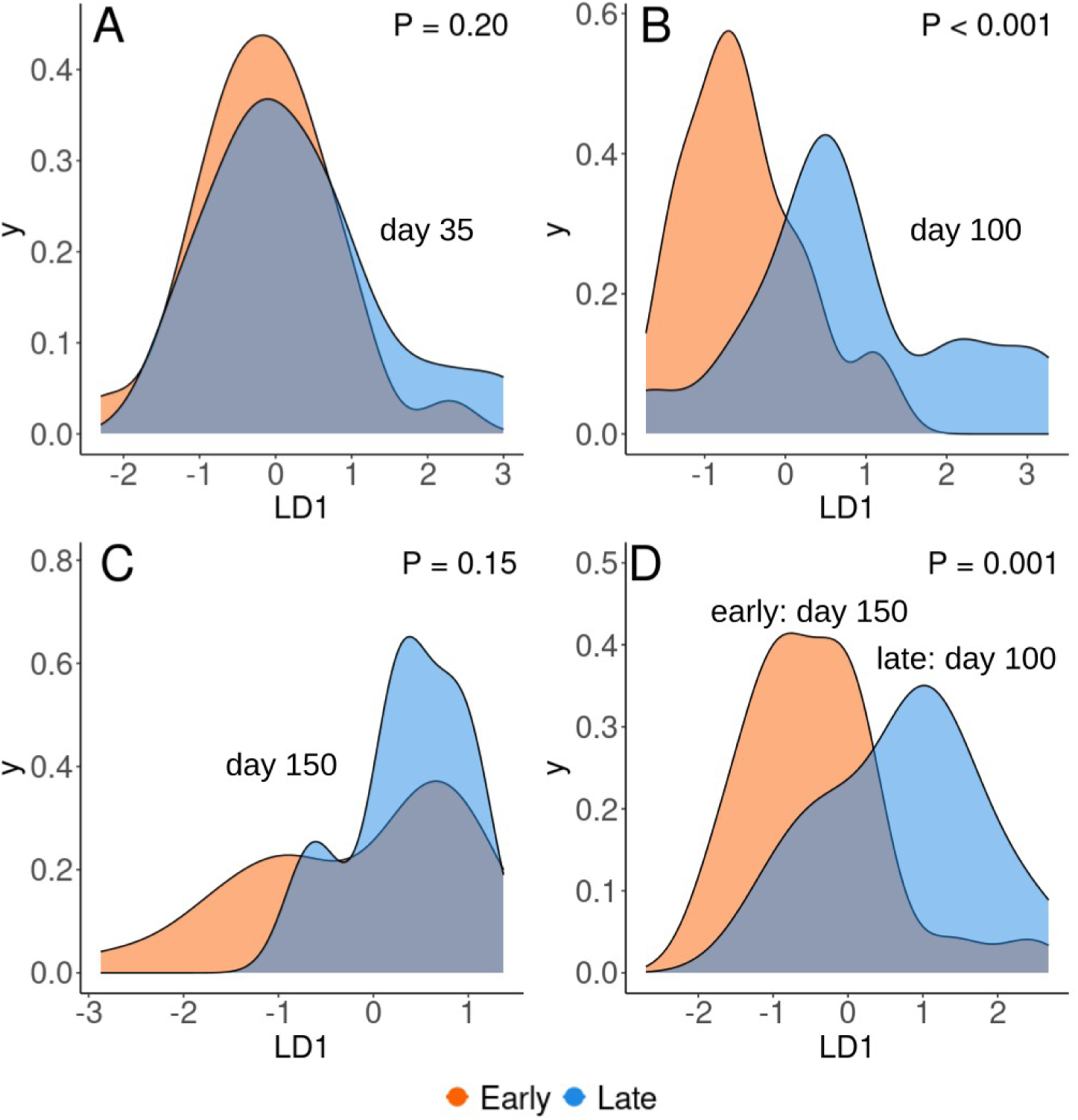
Results of the Linear Discriminant Analysis (LDA) testing the differences between early-*vs.* late-chicks according to RBC mitochondrial metabolic rates measured at 35 days (A), 100 days (B) and 150 days (C) post-hatching. Comparisons between chicks facing similar environmental conditions (early-chicks day 150 and late-chicks day 100) are also presented. **(D)**. The separation between early- and late-chicks was tested as a function of their RBC mitochondrial metabolic profile, including *ROUTINE, CI, CI+II* and *LEAK*. P-values from ANOVA are shown. The x-axis represents the first discriminant function (LD1), which is a linear combination of four variables: *ROUTINE, CI, CI+II, LEAK*. The y-axis shows the density distribution of both early- and late-chicks on this axis. In (A) and (C), the distributions of both groups overlap: as a result early- and late-chicks cannot be separated according to RBC mitochondrial metabolic rates tested. However, in (B) and (D), the overlap of the distributions is reduced and both groups are significantly distinct when taking into account their RBC mitochondrial metabolic profile.

### Predictions of growth trajectories by RBC mitochondrial metabolism

Except for *FCR_R/OL_*, differences in body mass gain from day 35 to day 100 were neither associated with RBC mitochondrial metabolic rates measured at day 35 or at day 100 (all P > 0.07, Table S1). We found a significant association between body mass gain and *FCR_R/OL_* measured at day 100 (P = 0.04), however this association was driven by a single measure from a late-chick with a high *FCR_R/OL_* (> 1) at day 100 (see Fig.S2A) and was not significant anymore if this value was excluded (P = 0.08). Body mass gain was strongly associated with the hatching group, with lower body mass gain for the late-chicks (all P < 0.001, Table S1).

Again, except for *FCR_R/OL_*, changes in body mass from day 100 to day 150 were not associated with RBC metabolic rates measured at day 100 or at day 150 (all P > 0.07, Table S2). For *FCR_R/OL_*, changes in body mass were significantly associated with *FCR_R/OL_* measured at day 150 (P = 0.01, Table S2).

## Discussion

We found marked differences in the growth trajectories of king penguin chicks that hatched early or late in the breeding season. Early-and late-chicks had similar body masses and sizes at 35 days post-hatching. In line with previous studies, chick body mass rapidly increased from 35 to 100 days post-hatching, and then decreased (significantly for the early-chicks) from 100 to 150 days post-hatching (Geiger et al., 2012; Stier et al., 2014). This decrease in body mass is most likely linked to a reduction in the number of feeding events for both groups following the rapid growth period (Saraux et al., 2012). Parental chick feeding decreases in association with the antarctic polar front position (abundant distribution of myctophid fishes), shifting southward during winter and therefore extending the foraging trips for the parents (Bost et al., 2004; Cherel & Ridoux, 1992; Freeman et al., 2016; Jouventin et al., 1994). Whereas the accurate position of the antarctic polar front varies between years, its withdrawal may even take place a few weeks earlier before the beginning of winter (starting in May) (Charrassin & Bost 2001; Cristofari et al., 2018; Freeman et al., 2016; Stonehouse 1956, 1960). For instance, Charrassin & Bost (2001) reported an average foraging trip distance of *ca* 340km, *ca* 900km and *ca* 1610km during austral summer, autumn and winter respectively for the king penguin population in Crozet (data collected between 1994-1997, but similar to results presented in Bost et al. 2015). In our study-sample, late-chicks were around 100 days-old when facing winter conditions, while early-chicks were around 150 days-old. This means that the core growth period (from day 35 to day 100) took place during favorable conditions for the early-chicks (summer-autumn), while the late-chicks core growth period occurred at the end of autumn-beginning of winter. When comparing similar ages, late-chicks were significantly lighter and smaller than the early-chicks from 100 days post-hatching and onwards as reported in the literature (Fernandes, 2023; Stier et al., 2014; VanHeezik et al., 1993). Prior studies showed that late-chicks can experience faster growth rates, and as a result express different markers of physiological stress (higher ROS levels and telomere loss) (Geiger et al., 2012; Stier et al., 2014). While we did not strictly compare the body masses of early and late-chicks a few days after hatching in our study, and therefore cannot assess if late-chicks experienced a faster growth rate at the beginning of the rearing period, we show that late-chicks did not manage to catch-up to an equivalent body mass and size of early-chicks before winter.

As expected, we found a significantly lower fledging success for the late-chicks, probably because of their small body mass and size, which prevented them from surviving during winter (Fernandes, 2023; Stier et al., 2014; VanHeezik et al., 1993; Weimerskirch et al., 1992). In our case, it is important to consider that we selected individuals that survived at least until 35 days to include them in this study - constituting a biased sample-size with selective disappearance (mortality before 35 days post-hatching not taken into account). However, similar patterns have been found on bigger sample-sizes (Fernandes, 2023).

A key purpose of this study was to test if early- and late-chicks expressed different red blood cell (RBCs) mitochondrial metabolic rates during the core growth period (from day 35 to day 100), but also during the winter period. We expected late-chicks to have higher RBC mitochondrial metabolism and a higher coupling efficiency to meet the energy requirement linked to a faster growth rate. As late-chicks can express higher oxidative stress (Stier et al., 2014), we could expect a higher respiration of complex I (*CI*) and proton leak (*LEAK*) for the late-chicks. Both the oxygen consumption before RBC permeabilization (endogenous respiration, *ROUTINE*) and the ratio between the endogenous respiration and the maximum respiration of complexes I and II (*FCR_R/OL_*) were higher for the late-chicks at 100 days post-hatching. In other words, at similar ages, late-chicks consumed more oxygen and had a higher usage of the complexes I and II respiration maximal capacity at the end of the core growth period (day 100). Additionally, the maximal respiration capacity of the complex I (*CI*) was significantly higher in the late-chicks at all ages. Yet, the respiration linked to oxidative phosphorylation did not differ between early- and late-chicks, meaning that late-chicks had higher, but not more efficient RBC metabolism than the early-chicks when comparing same ages (i.e. no difference observed in *OXPHOS* coupling efficiency).

These results were also confirmed by linear discriminant analysis (LDA), which was able to well separate the early- and late-group when taking into account *ROUTINE, CI, CI+II* and *LEAK*. Interestingly, these metabolic differences at day 100 were not associated with the discrepancy in chick body mass between early- and late-chicks, but did correlate with the date of sampling: a proxy of the environmental conditions. Our results seem reasonable in light with the fact early-chicks reached 100 days-old during austral-autumn (*ca* end of April) and the late-chicks during winter (*ca* beginning of June). Different RBC metabolic profiles can be linked to different environmental conditions, but also with the stress associated with growing in adverse conditions and experiencing food restriction during growth (Blas, 2015). This hypothesis would be supported by the fact that late-chicks express higher corticosterone levels - hormone involved in *stress response* (Blas, 2015; Stier et al., 2014, but see Breuner et al., 2012; Cockrem 2013).

As suggested above, late-chicks may express a higher RBC mitochondrial metabolism (higher endogenous respiration, higher respiration of the complex I, and higher respiration of the complexes I and II) to ensure a higher production of ATP to sustain the energy requirements linked to a faster growth rate. This hypothesis would be supported by recent research reporting that late-born king penguin chicks may have some adaptations linked to a faster growth, by the over-expression of genes potentially involved in food uptake efficiency, body mass increase, and accumulation of abdominal fat (Fernandes, 2023).

An alternative hypothesis is the differences in fasting stages between chicks. Indeed, since late-chicks reached 100 days in the beginning of winter (*vs.* autumn for early-chicks), they may experience more frequent and longer fasting events than early-chicks at the same age. Previous studies carried out in king penguin chicks reported a variation in skeletal muscle mitochondrial metabolism according to fasting stages (increase in metabolism efficiency between *phase II* and *phase III* of fasting) (Bourguignon et al., 2017; Monternier et al., 2014; Teulier et al., 2013). Here, a higher complex I respiration in the late-chicks could be linked to a variation in the usage of metabolic substrates compared to early-ones and could reflect energetic adjustments linked to nutritional status (last feeding event), substrates availability and body composition (lipid and protein stores), but also fasting stage (glycolysis with NADH generation *vs*. β-oxidation with production of FADH_2_-derived electrons) (Bernard et al., 2003; Bochkova et al., 2025; Liesa & Shirihai, 2013; Secor & Carey, 2016). For instance, RBC mitochondrial metabolism has been shown to vary (to some extent) according to fasting stages in adult king penguins (Cossin-Sevrin et al., 2025). Thus, differences in RBC mitochondrial metabolic rates could be also associated with different fasting stages between early- and late-chicks.

All RBC mitochondrial metabolic rates decreased with age, as reported in other species (Cossin-Sevrin et al., 2022, 2023; Stier et al., 2022). In these studies, such decrease in RBC mitochondrial metabolism was closely linked to a reduction in RBC mitochondrial content (mitochondrial density) across the growth period and we may expect similar patterns during the core growth period in king penguin chicks (day 35 to day 100). A large reduction in mitochondrial content per cell would lead to a decrease in RBC mitochondrial metabolism, providing an explanation for our results. In our case, as body masses and sizes differed between early- and late-chicks at day 100 days, one possibility is that early- and late-chicks were not at similar developmental stages when drawing comparisons at similar ages. Distinct developmental stages would explain different RBC mitochondrial metabolic rates (e.g. through a variation in mitochondrial content as explained above). Additionally, rapid growth has been shown to increase mitochondrial content per cell (in japanese quails), which may also explain higher metabolic rates for late-chicks (Jimenez et al., 2014). Yet, if the late-chicks possessed a higher number of mitochondria per cell, we could expect all RBC mitochondrial metabolic rates to increase (including *OXPHOS_CI+II_*, and *CI+II*).

Most of the RBC mitochondrial metabolic rates were affected by sampling year. This effect is most likely due to a bias in our sample-size, with very few late-chicks monitored during the season 2021-2022, 2022 being an overall poor-quality year with lower chick-survival at the level of the colony (only 2 late chicks monitored from 100 post-hatching for this season). However, mitochondrial metabolism in endotherms has been shown to vary according to environmental conditions: including ambient temperature and food restrictions (Gyllenhammer et al., 2020; Le Roy et al., 2021; Mahalingam et al., 2020; Zitkovsky et al., 2021). Here, we cannot totally exclude the impact of different environmental conditions between years and further studies would be needed to comprehensively assess the effect of the environment on king penguins chick RBC mitochondrial metabolism.

One of our aims was to assess if differences in growth trajectories between early- and late-chicks could be predicted by differences in RBC mitochondrial metabolism. We only found a positive association between changes in body mass after the core growth period (day 150 – day 100) and *FCR_R/OL_* measured at day 150. This would suggest that individuals expressed a lower usage of their maximal respiration of complexes I and II when experiencing a body mass drop between day 100 and day 150. However, R^2^ for this set of analysis was low (R^2^ = 0.3), meaning that differences in RBC mitochondrial metabolism were not the main contributor creating differences in growth. As suggested above, differences in environmental conditions and in parental food provisioning (and stress associated) were probably the main contributors explaining differences in RBC metabolic profiles at day 100 between early and late-chicks. Furthermore, this hypothesis is strengthened by the fact that RBC mitochondrial metabolic profile were similar between early- and late-chicks at day 150, when both groups of chicks were facing winter conditions. It is worth noting that when comparing chicks at different ages during a similar period (early-chicks at day 150 and late-chicks at day 100, see Fig.1), RBC metabolic profiles remained different between the hatching groups (early *vs*. late), suggesting that other factors influenced differences in chick metabolism (as suggested above: stress, faster growth rate, mitochondrial content per cell).

In conclusion, phenotypic differences between early- and late-born king penguin chicks were visible from 100 days post-hatching and onwards. Late-chicks had lower body mass and size, most likely explaining low survival chances (fledging success) compared to early-chicks. When comparing similar ages, late-chicks had higher endogenous RBC mitochondrial metabolism (*ROUTINE*), higher usage of the complex I respiration (*CI*), but also higher usage of the complexes I and II respiration (*CI+II*). These differences were more pronounced 100 days post-hatching, most likely due to differences in environmental conditions: number of feeding events and/or the stress associated with facing adverse conditions at younger age for the late-chicks. For instance, adverse weather events have been shown to impact king penguin chick metabolic budget and heat production (heterothermy), most probably impacting the animal metabolic rate (Eichhorn et al., 2011). Due to global changes, the breeding conditions are expected to change for king penguin population in Crozet archipelago as the polar front contracts around the Antarctic continent moving further from the breeding colony, and as sea surface temperatures modify the growth and size of king penguin prey (Brisson-Curadeau et al., 2022; Péron et al., 2012; Saunders et al., 2020). These scenarios are expected to impact the king penguins breeding success. Understanding how nutritional changes (quantity and quality) may impact the king penguin chicks’ metabolism, growth and survival (both early and late-born chicks) would ultimately help in predicting the response of king penguin populations facing changing conditions.

## Acknowledgments

This research was supported by the French Polar Research Institute (IPEV; project 119 ECONERGY), by the Centre National de la Recherche Scientifique (CNRS) and the Zone Atelier Antarctique (ZATA). We thank the Terres Autstrales et Antarctiques Françaises (TAAF) for their logistic support on the field. We are grateful to Antoine Stier for his advice on mitochondrial measurements, his initial work on mitochondria in freely breeding king penguins, and his contribution to the 119 project. We thank the IPEV projects 137, 131 and 394 for their help on the field. N.C-S was supported by Maupertuis Grant; the Biology, Geography and Geology doctoral program of the University of Turku and the Alfred Kordelin Foundation (Alfred Kordelinin Säätio) at the time of writing. M.L, C.B, M.F, T.F, N.G, A.C were funded by the IPEV as Civil Service Volunteers. The ECONERGY king penguin project is part of the long-term Studies in Ecology and Evolution (SEE-Life) program of CNRS.

## Ethics

All the procedures were approved by the French Ethical Committee (APAFIS#31268-2021042117037897 v3 and APAFIS#16465-2018080111195526 v4) and the Terres Australes et Antarctiques Françaises (Arrêtés TAAF A-2020-82 and A-2021-49).

## Authors contribution

V-A.V, J-P.R and P.B designed the study. M.L, C.B, C.L, N.C-S, T.F, M.F, A.C and V-A.V conducted the fieldwork and collected the samples. M.L, C.B, N.C-S and N.G conducted the mitochondrial respirometry measurements. N.C-S performed statistical analysis under the supervision of K.A and S.R. N.C-S wrote the first version of the manuscript under the supervision of K.A and S.R. All co-authors revised the manuscript.

## Supplementary Materials

**Table S1:** Results of linear models (LMs) testing the effect of the chick body mass during the whole growth period (from day 35 to day 150 post-hatching) on RBC mitochondrial metabolic rates (upper part) and the effect of the date of sampling (in the number of weeks in the year, proxy of environmental conditions) on chick RBC mitochondrial metabolic rates measured at day 100 (bottom part). Estimates are reported with their standard error. Bold indicates significance (P < 0.05). Because of the correlation between the date of sampling and the early-late group, we only tested the impact of the date on RBC mitochondrial metabolic rates measured at day 100, when the differences between early- and late-chicks were significant (see Results).

| Predictors | *ROUTINE* |  | *CI* |  | *FCR<sub>R/OL</sub>* |  |
| --- | --- | --- | --- | --- | --- | --- |
| | Estimate ± SE | P | Estimate ± SE | P | Estimate ± SE | P |
| (Intercept) | 6.05 ± 0.30 | **< 0.001** | 6.33 ± 0.37 | **< 0.001** | 0.48 ± 0.03 | **< 0.001** |
| mass (kg) | -0.07 ± 0.07 | 0.35 | -0.03 ± 0.09 | 0.76 | -0.004 ± 0.007 | 0.58 |
| age day 100 | -1.83 ± 0.39 | **< 0.001** | -1.72 ± 0.48 | **< 0.001** | -0.01 ± 0.04 | 0.73 |
| age day 150 | -2.18 ± 0.37 | **< 0.001** | -2.16 ± 0.46 | **< 0.001** | 0.007 ± 0.04 | 0.84 |
| year (2021) | 0.41 ± 0.23 | 0.08 | -0.40 ± 0.29 | 0.18 | 0.05 ± 0.02 | **0.05** |
| Random effects |  |  |  |  |  |  |
| $\sigma^2$ | 1.18 | | 1.43 | | 0.11 | |
| $\tau_{00}$ bird | 0.23 | | 0.45 | | 0.05 | |
| n observations | 122 |  | 123 |  | 122 |  |
| N individuals | 55 |  | 55 |  | 55 |  |
| Marginal R<sup>2</sup> /conditional R<sup>2</sup> | 0.51 / 0.49 |  | 0.38 / 0.32 |  | 0.21 / 0.07 |  |
| (Intercept) | -1.19 ± 1.50 | 0.43 | 1.04 ± 1.50 | 0.49 | 0.003 ± 0.16 | 0.99 |
| Number of weeks | 0.25 ± 0.07 | **0.001** | 0.16 ± 0.07 | **0.03** | 0.02 ± 0.01 | **0.008** |
| year (2021) | 0.76 ± 0.49 | 0.13 | 0.25 ± 0.49 | 0.61 | 0.07 ± 0.05 | 0.24 |
| n observations | 39 |  | 39 |  | 39 |  |
| N individuals | 39 |  | 39 |  | 39 |  |
| R<sup>2</sup> | 0.25 |  | 0.13 |  | 0.18 |  |

**Fig.S1:**
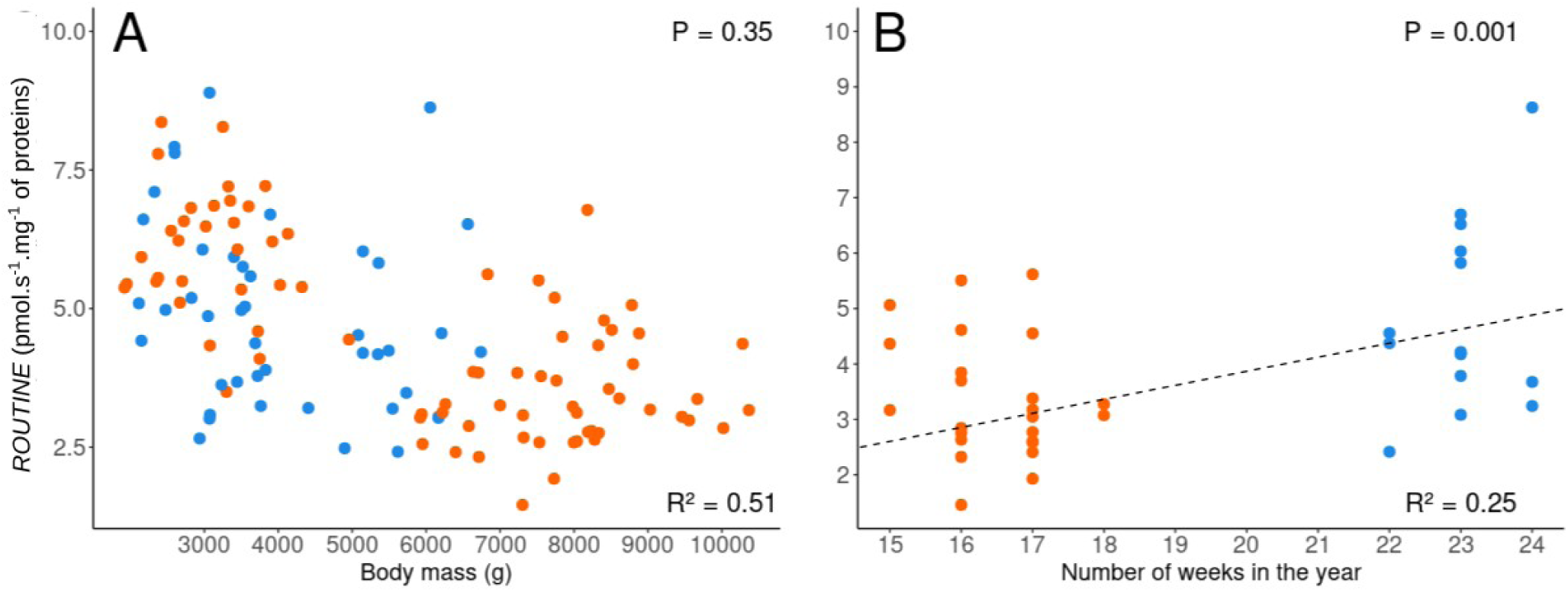
Associations between the chick body mass (g) during the whole growth period (from day 35 to day 150 post-hatching) (A); and the date of sampling (in weeks) (B) with *ROUTINE* metabolic rate. Raw data are plotted. Results presented from Linear mixed models can be found in Table S1.

**Table S2:** Results of linear models (LMs) testing the effect of RBC mitochondrial metabolic rates (measured at 35 and day 100 post-hatching), the hatching group (early *vs.* late) and the year (2020 *vs.* 2021) on chick body mass gain between during the core growth period from day 35 to day 100 post-hatching. Estimates are reported with their standard error. Bold indicates significance (P<0.05).

| Predictors | ROUTINE |  |  | CI |  |  | CI+II |  |  | LEAK |  |  |
| --- | --- | --- | --- | --- | --- | --- | --- | --- | --- | --- | --- | --- |
|  | Estimate | SE | P | Estimate | SE | P | Estimate | SE | P | Estimate | SE | P |
| (Intercept) | 4951.1 | 1192.9 | < 0.001 | 3851.6 | 1420.6 | 0.01 | 2725.3 | 1880.9 | 0.16 | 3522.6 | 952.1 | 0.001 |
| metabolic rate (day 35) | -130.4 | 197.4 | 0.51 | 113.4 | 175.9 | 0.53 | 155.7 | 105.5 | 0.15 | 957.4 | 504.3 | 0.07 |
| metabolic rate (day 100) | 257.7 | 159.6 | 0.12 | 140.3 | 167.2 | 0.41 | 47.2 | 114.6 | 0.68 | -175.5 | 329.9 | 0.60 |
| hatching group (late) | -3737.0 | 602.9 | < 0.001 | -3532.2 | 553.8 | < 0.001 | -3547.3 | 521.0 | < 0.001 | -3506.2 | 525.9 | < 0.001 |
| year (2021) | -76.2 | 500.3 | 0.88 | -31.7 | 465.3 | 0.95 | -43.9 | 424.3 | 0.92 | -745.4 | 635.2 | 0.25 |
| n observations | 30 |  |  | 30 |  |  | 30 |  |  | 30 |  |  |
| Multiple R<sup>2</sup> | 0.67 |  |  | 0.65 |  |  | 0.66 |  |  | 0.68 |  |  |

| Predictors | OXPHOS coupling efficiency |  |  | FCR<sub>R/OL</sub> |  |  |
| --- | --- | --- | --- | --- | --- | --- |
|  | Estimate | SE | P | Estimate | SE | P |
| (Intercept) | 8590.2 | 5838.4 | 0.154 | 5211.8 | 873.6 | < 0.001 |
| metabolic rate (day 35) | -4883.1 | 6836.3 | 0.482 | -3141.3 | 1741.3 | 0.08 |
| metabolic rate (day 100) | 750.7 | 3163.5 | 0.814 | 3081.2 | 1387.5 | 0.04 |
| hatching group (late) | -3296.8 | 537.5 | < 0.001 | -3957.2 | 540.5 | < 0.001 |
| year (2021) | -413.2 | 728.5 | 0.576 | 208.7 | 456.6 | 0.65 |
| n observations | 30 |  |  | 30 |  |  |
| Multiple R<sup>2</sup> | 0.64 |  |  | 0.71 |  |  |

**Table S3:** Results of linear models (LMs) testing the effect of RBC mitochondrial metabolic rates (measured at 100 and day 150 post-hatching), the hatching group (early *vs.* late) and the year (2020 *vs.* 2021) on changes in chick body mass after the core growth period from day 100 to day 150 post-hatching. Estimates are reported with their standard error. Bold indicates significance (P<0.05).

| Predictors | *ROUTINE* |  |  | *CI* |  |  | *CI+II* |  |  | *LEAK* |  |  |
| --- | --- | --- | --- | --- | --- | --- | --- | --- | --- | --- | --- | --- |
|  | Estimate | SE | P | Estimate | SE | P | Estimate | SE | P | Estimate | SE | P |
| (Intercept) | -730.8 | 817.8 | 0.38 | -245.3 | 711.3 | 0.73 | -111.5 | 1022.4 | 0.91 | -499.0 | 718.1 | 0.49 |
| metabolic rate (day 100) | -258.3 | 155.5 | 0.11 | -222.9 | 134.7 | 0.11 | -102.7 | 102.9 | 0.33 | -44.7 | 285.9 | 0.88 |
| metabolic rate (day 150) | 295.2 | 180.9 | 0.12 | 156.3 | 131.6 | 0.25 | 63.8 | 122.1 | 0.61 | 115.1 | 472.5 | 0.81 |
| hatching group (late) | 276.8 | 412.7 | 0.51 | 147.5 | 399.7 | 0.72 | 15.0 | 401.3 | 0.97 | -69.5 | 420.7 | 0.87 |
| year (2021) | -378.8 | 344.3 | 0.28 | -387.1 | 352.3 | 0.28 | -428.3 | 375.2 | 0.26 | -565.9 | 640.4 | 0.39 |
| n observations | 31 |  |  | 31 |  |  | 31 |  |  | 31 |  |  |
| Multiple R<sup>2</sup> | 0.23 |  |  | 0.23 |  |  | 0.11 |  |  | 0.08 |  |  |

| Predictors | *OXPHOS coupling efficiency* |  |  | *FCR<sub>R/OL</sub>* |  |  |
| --- | --- | --- | --- | --- | --- | --- |
|  | Estimate | SE | P | Estimate | SE | P |
| (Intercept) | 1316.2 | 3745.2 | 0.73 | -1662.7 | 928.3 | 0.09 |
| metabolic rate (day 100) | -479.0 | 2518.2 | 0.85 | -2890.4 | 1543.6 | 0.07 |
| metabolic rate (day 150) | -1539.4 | 3944.9 | 0.70 | 5262.9 | 1947.5 | **0.01** |
| group (late) | -118.5 | 403.2 | 0.77 | 182.1 | 368.7 | 0.63 |
| year (2021) | -796.0 | 703.2 | 0.27 | -765.7 | 349.0 | **0.04** |
| n observations | 31 |  |  |  | 31 |  |
| Multiple R<sup>2</sup> | 0.08 |  |  |  | 0.30 |  |

**Fig.S2:**
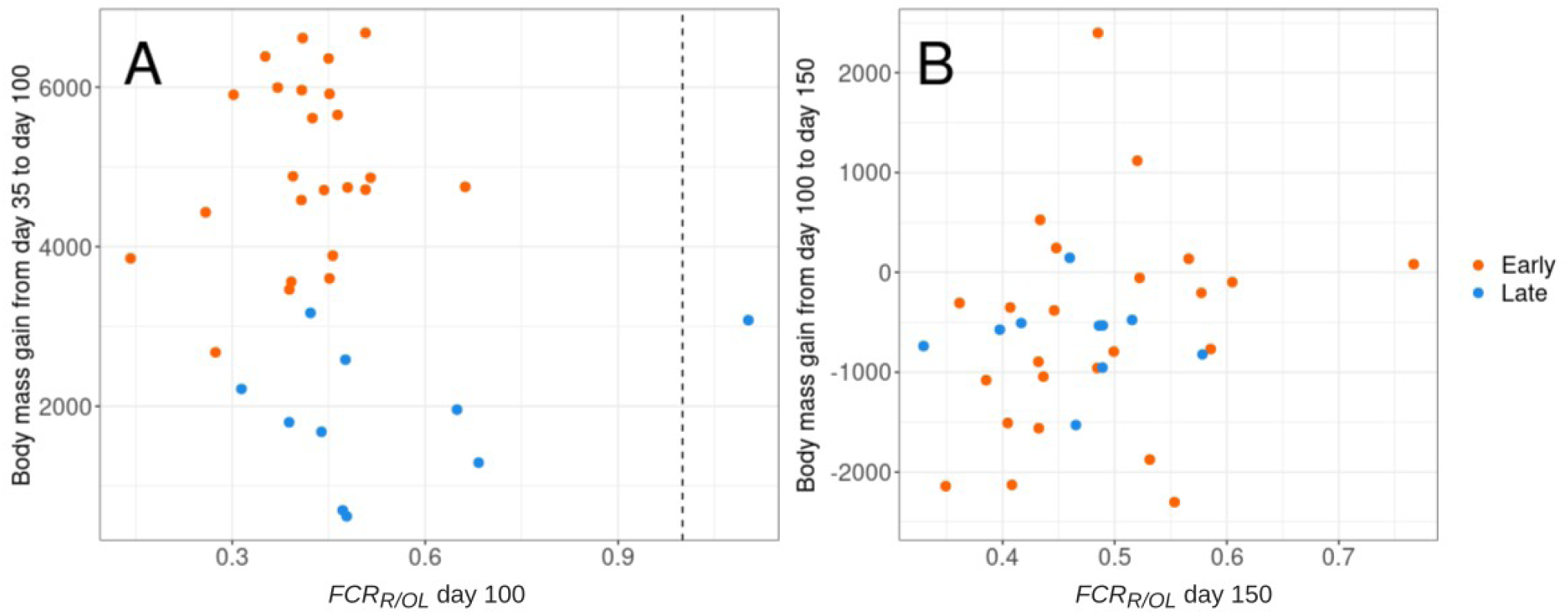
Association between the body mass gain (g) during the core growth period (from day 35 to day 100 post-hatching) and *FCR_R/OL_* measured at day 100 (A); and changes in body mass after the core growth period (from day 100 to day 150 post-hatching) and *FCR_R/OL_* measured at day 150 (B). Raw data are plotted, For (A) the association did not remain significant when the highest value for *FCR_R/OL_* was removed from the analysis (value on the right of the dashed line). The average (± SD) body mass gain for early- and late-chicks were respectively 4928.5g ± 1136.9 and 1949.1g ± 852.9 between 35 and 100 days post-hatching. Body mass loss between 100 and 150 days post-hatching were -636.9g ± 1099.8 for the early-chicks and -7796g ± 583.8 for the late-chicks.

